# Min-frame transformation enables more sensitive viral genome alignment

**DOI:** 10.64898/2026.05.20.726535

**Authors:** Ryan D. Doughty, Ankhi Banerjee, Bryce Kille, Tandy Warnow, Todd J. Treangen

## Abstract

**Motivation:** Maximal unique matches (MUMs) are a fundamental primitive in genome comparison, where they serve as high-confidence anchors for downstream multiple genome alignment. However, because MUMs rely on exact string matching, their effectiveness degrades with increased genome divergence and larger sets of genomes, inhibiting their ability to recover long homologous regions and reducing the number of base pairs covered by the multiple genome alignment. Additionally, existing approaches that improve robustness to mutation, such as spaced seeds or translated alignment methods, introduce trade-offs in specificity, scalability, or computational complexity.

**Methods:** To address this gap, we introduce the Min-Frame Transformation (MFT), a deterministic encoding of nucleotide sequences to sequences over a transformed alphabet that preserves the coordinate structure of the original sequence. At each position, the MFT selects a *k*-mer from a local window according to a fixed global ordering and assigns it a character in the transformed alphabet via a predefined mapping. This process captures local sequence context and can mask the impact of mutations, increasing the likelihood that homologous regions remain detectable as exact matches. The resulting transformed sequences can be indexed using standard string data structures, such as suffix arrays and suffix trees, enabling efficient extraction of MUMs without modifying existing algorithms.

**Impact:** The MFT is a novel computational approach for improving the robustness of MUM-based seeding for genome alignment by producing longer and more contiguous matches that span a greater fraction of the genome, leading to improved alignment coverage and SNP recall. Altogether, these improvements have the potential to result in improvements for downstream viral genome analysis applications such as phylogenetic inference and transmission analysis.

**Funding:** *Tandy Warnow*: NSF grant 2316233

*Todd J. Treangen*: NSF grants 2126387, 2239114, NIH grants U19-AI144297, P01-AI152999

## 1 Introduction

Given a set of DNA sequences, sequence similarity searches to identify homologous regions are the first and often most critical step in comparative genomic analyses [1, 2]. Among these seeding strategies are maximal unique matches (MUMs). By definition, a MUM is a sub-string that occurs exactly once in two genomes and cannot be extended in either direction without encountering a mismatch [3, 4]. MUMs occupy a unique niche among possible alignment seeds due to several desirable properties. *i)* Because a MUM must be unique in every input genome, it cannot anchor within repetitive regions, unlike maximal exact matches (MEMs) [5] or *k*-mer based approaches. This eliminates the need for post hoc filtering of repeat-induced matches. *ii)* Long, exact unique matches between two large genomes are highly unlikely to appear by chance. Thus, sufficiently long MUMs that occur across the input set represent high-confidence signal of potential homologous regions that can be used to anchor downstream alignments [6, 7]. *iii)* Efficient algorithms based on suffix trees, suffix arrays, and FM-index variants enable scalable identification of MUMs across large datasets [7, 8, 9, 10, 11]. As a result of these three factors, MUMs, and their multi-genome generalization multi-MUMs, form the foundation of many widely used whole-genome alignment tools for closely related microbial genomes [3, 6, 7, 12].

However, all of these approaches suffer from the limitation that multi-MUMs dissolve with increasing sequence divergence and the number of genomes. Because these seeds rely on perfect string matches, a single nucleotide polymorphism (SNP), insertion, or deletion within one genome can break a potential anchor, significantly reducing the length, density, and genome coverage of the resulting seed set. This effect becomes even more pronounced in alignments involving many genomes or highly divergent taxa, where increasing sequence variation and sample size substantially reduce the number and length of perfect maximal matches shared across all genomes [2].

Several approaches have been developed to address this sensitivity issue, but each introduces trade-offs. Flexible seeding strategies such as spaced seeds [12, 13, 14, 15] can improve tolerance to mutations but often reduce specificity or increase chaining complexity. Alternatively, amino acid–based approaches such as promer [8] translate genomes and search for matches in protein space to exploit higher conservation [16]. While often more effective for detecting distant homology [17], these methods introduce substantial computational and algorithmic overhead due to translation and the need to consider all six reading frames. This leads to ambiguous and overlapping matches that can be difficult to resolve and slower to compute. Additionally, no existing software can compute multi-MUMs in amino acid space across input sets with more than two genomes.

To overcome the susceptibility of nucleotide-based multi-MUMs to sequence variation without losing the computational simplicity, we introduce the Min-Frame Transformation (MFT). The MFT is a deterministic mapping that encodes genomic sequences into equivalent-length sequences over a separate alphabet. Similar to minimizer schemes [18], the MFT first *O* selects a *k*-mer from each overlapping window of length *w* using a global *k*-mer ordering _*k*_. A mapping *M* then assigns each selected *k*-mer to an ASCII character in the transformed alphabet, producing the final encoded sequence. By selecting *k*-mers from the surrounding context and mapping them to a separate alphabet, the MFT effectively enables local masking of mutations, which allows the genomes to be better suited for downstream multi-MUM inference. Importantly, the MFT emits a coordinate-preserving transformed string, making it directly compatible with optimized full-text MUM search algorithms, provided the implementation is not restricted to a nucleotide alphabet. All in all, MFT sequences produce more sensitive anchors while retaining the simplicity of nucleotide MUM search, bridging the gap between nucleotide precision and protein-level recall.

The primary criterion when evaluating multi-MUMs is their ability to support the creation of long, contiguous multiple alignments that cover the genetic variation of the genomes. We show that multi-MUMs identified on transformed genomes consistently yield longer alignments than those derived from the original nucleotide sequences, while also retaining more SNP information that could be useful for downstream analyses such as phylogenetic inference and transmission reconstruction. Through the measurement of multi-MUM coverage, we also show these alignment gains primarily stem from the fact that the transformations produce longer and more contiguous multi-MUMs that cover more of the genomes.

We also provide a fully functional toolkit called mft-tools to assist others to define, evaluate, and optimize different *k*-mer mappings and orderings, as well as transform their own DNA sequences. The code is publicly available at https://github.com/treangenlab/mft-tools.

## 2 Notation and Preliminaries

Here we introduce notation and other preliminaries referenced throughout this work.

### Notation

We use standard string notation throughout. Let Σ_*σ*_ denote an alphabet of *σ* characters. In particular, the nucleotide alphabet is Σ_*n*_ = {*A, C, G, T}*. A sequence 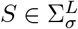 is a string of length *L* over Σ_*σ*_, and *S*[*i* : *j*] denotes the substring from position *i* to *j* (inclusive). A *k*-mer is a string of length *k*, and 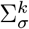 denotes the set of all such strings over a particular alphabet. For a given position *i*, we define a window *W*_*i*_ = *S*[*i* : *i* + *w*− 1] of length *w*, which induces a set of overlapping *k*-mers 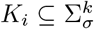 with *k ≤w*. Throughout, *k* denotes the *k*-mer length and *w* the window length. Note that *w* refers to the nucleotide span of the window, rather than the number of *k*-mers it contains.

#### ▸Definition 1.

***(Order)** An order 𝒪_k_ on k-mers is a function 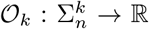,such that 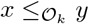 if and only if 𝒪_k_(x) ≤ 𝒪_k_(y).*

While many orderings in classical minimizers schemes are based on hash values or functions, non-random orderings will be explored and are a particular point of emphasis throughout this work.

#### ▸Definition 2.

***(Mapping)** A mapping M is a function 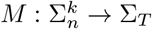 that assigns each k-mer in 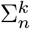to a character in a separate transformed alphabet Σ_T_*.

### Sampling function terminology

Given the similarity of the Min-Frame Transformation to *k*-mer sampling schemes, we also introduce relevant sampling terminology, particularly for minimizers. We follow the definitions of [19].

#### ▸Definition 3.

***(Local Schemes)** A local scheme is a function f : Σ ^w^→[w] that, given a window W, selects the k-mer starting at position f (W) in W. That is, the sampled k-mer is W [f (W) : f (W) + k − 1]*.

#### ▸Definition 4.

***(Forward Schemes)** A forward scheme f is a local scheme when for any two consecutive windows W and W^′^ it holds that f (W) ≤ f (W^′^) + 1*.

#### ▸Definition 5.

***(Minimizers)** A minimizer is a forward scheme defined by a total order 𝒪_k_ on k-mers and selects the leftmost minimal k-mer in a window W*

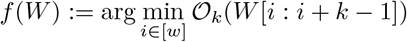

#### Maximal Unique Matches

We now define terminology relevant to maximal unique matches and its variations.

##### ▸Definition 6.

***(Unique Substring)** Let S ∈* Σ and P ∈ Σ^𝓁^. We say P is unique in S if there exists exactly one index i such that S[i : i + 𝓁 − 1] = P*.

##### ▸Definition 7.

***(Maximal Match)** Let S*_1_, *S*_2_ ∈ Σ* *and P* ∈ Σ^𝓁^ *such that S*_1_[*i* : *i* + 𝓁 − 1] = *S*_2_[*j* : *j* + 𝓁 − 1] = *P*. *We say P is maximal if it cannot be extended in either direction, i.e., S*_1_[*i* − 1] ≠ *S*_2_[*j* − 1] *and S*_1_[*i* + 𝓁] ≠ *S*_2_[*j* + 𝓁].

##### ▸Definition 8.

***(MUM)** Let S*_1_, *S*_2_ *∈* Σ*. *A substring P∈* Σ^𝓁^*is a maximal unique match (MUM) if it occurs in both S*_1_ *and S*_2_, *is unique in each sequence, and is maximal with respect to S*_1_ *and S*_2_.

##### ▸Definition 9.

***(Multi-MUM)** Let* 𝒮 = *{S*_1_, …, *S*_*n*_*} be a set of sequences with S*_*i*_ ∈ Σ*. *A substring P ∈* Σ^𝓁^*is a multi-MUM if it occurs in every sequence in* 𝒮, *is unique in each S*_*i*_, *and is maximal across the sequences in* 𝒮.

##### ▸Definition 10.

***(Subset Multi-MUM)** Let* 𝒮 = *{S*_1_, …, *S*_*n*_*} be a set of sequences with S*_*i*_ *∈* Σ*. *A substring P ∈* Σ^𝓁^*is a subset multi-MUM if there exists a subset* 𝒮*⊆* 𝒮 *with* 2 *≤* |𝒮′| = *k ≤ n such that P occurs exactly once in each sequence S*_*i*_ *∈ 𝒮′ and is maximal with respect to all sequences in* 𝒮′.

Although our definition of subset multi-MUMs is analogous to partial multi-MUMs used elsewhere [20], we find the term subset multi-MUM more precise and descriptive in this context. We show an in-depth schematic comparison of multi-MEMs vs multi-MUMs in **Appendix Figure 1**

**Figure 1.**
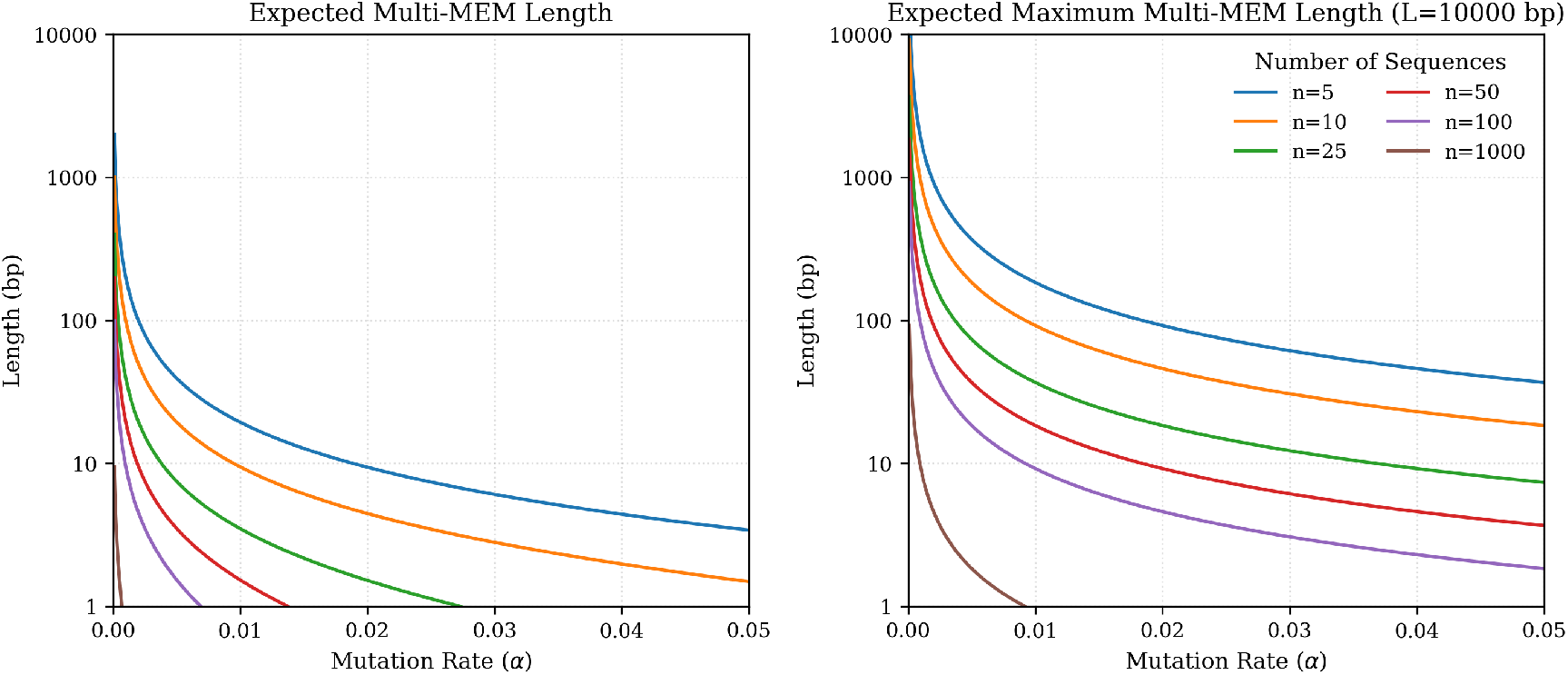
Expected Multi-MEM Length and Expected Maximum Multi-MEM Length of randomly mutated sequences as a function of the mutation rate *α* and the number of sequences *n*. Multi-MEMs quickly degrade in expected length as the mutation rate and number of sequences increase.

### 2.1 Expectation of multi-MEM Length for Random Genomes

In this section, we derive the expected length and expected maximum length of multiple maximal exact matches (multi-MEMs) for *n* sequences. We model the ancestral sequence *S* as an infinite stationary process and each descendant as evolving via independent and identically distributed (i.i.d.) Bernoulli trials with a uniform per-site mutation rate *α*. While real genomes exhibit non-random patterns and have highly conserved regions, this simplified model provides a mathematical expectation for multi-MEM presence under the assumption of random mutations. Additionally, since multi-MUMs are a strict subset of multi-MEMs, these results also serve as a conservative upper bound for multi-MUM conservation.

We assume a match at a specific position occurs if and only if the ancestral base is conserved across all *n* descendants, neglecting the negligible probability of convergent evolution. Under these assumptions, the probability of conservation for a single site in one sequence is 1− *α*. Thus, the probability *p* of a global match across *n* sequences at any given position is:

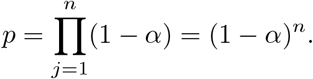

Let the random variable *X* represent the length of a contiguous match across *n* sequences. This follows a geometric distribution, where a match of length *x* corresponds to *x* successes of probability *p* followed by one failure. The probability mass function is:

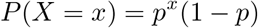

The expected match length *E*[*X*] is derived from the expectation of this distribution:

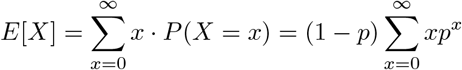

Then, using the identity for an arithmetico-geometric series 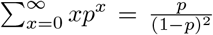 expression simplifies to:

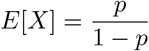

Finally, substituting *p* = (1− *α*)^*n*^ yields the exact expected length of a contiguous match across *n* sequences given a mutation rate *α*:

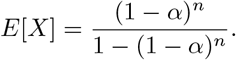

While this defines the expected length of a match, we often are also interested in the expected maximum match length within the sequences. To find the expected maximum match length *E*[*X*_*max*_] within sequences of finite length *L*, we consider the *L* possible starting positions. While these positions are not strictly independent, we apply the standard approximation for the distribution of the longest run of successes in Bernoulli trials [21, 22]. The probability that no match reaches a length *k* is:

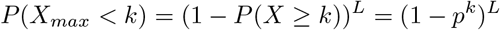

Given *p*^*k*^ is expected to be very small, we can use the approximation (1 −*x*)^L^ ≈*e*^−*Lx*^ for small *x*:

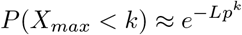

The expected maximum match length occurs approximately when the expected number of runs of length *k* within the genome is one, i.e. *Lp*^*k*^ ≈ 1. Intuitively, this corresponds to the transition point where a match of length *k* becomes unlikely to occur across all L starting points. Solving for *k*, we get:

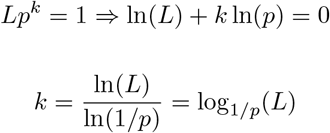

Substituting back in *p* = (1 *− α*)^*n*^:

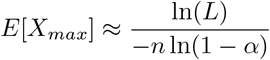

Finally, applying the Taylor expansion ln(1−*α*) *≈*−*α*, we get the final expected maximum length of an exact match for sequences of length L:

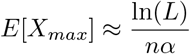

In **Figure 1**, we plot the expected match length and the approximate expected longest match length. Even for relatively small values of *n* and *α*, the expected match lengths degrade rapidly. For instance, with a mutation rate of 0.01 across 10 sequences, the expected match length is 10bp and the expected maximum match length is 100bp (assuming a 10,000bp sequence). However, increasing the mutation rate to 0.03 reduces these values to approximately 5 and 20, respectively, and increasing *n* to 50 further reduces the expected match length to nearly 2bp. These results illustrate the fragility of exact matches under uniformly random mutation processes and higher sequence counts.

Note that real genomes do not evolve in such a purely random manner. Biological sequences contain conserved regions shaped by functional constraints and selective pressures, and consequently MEMs and MUMs can still be identified across thousands of genomes within the same species [7, 23, 24]. Therefore, this analysis should primarily be interpreted as a motivating theoretical example illustrating how exact-match statistics deteriorate under increasing sequence variation, rather than as a realistic model of genome evolution.

## 3 Min-Frame Transformation

Motivated by the observation that the length and number of multi-MUMs can decrease substantially with increasing genome divergence and sample size, we propose the Min-Frame Transformation as an approach for reducing multi-MUM dropout in such settings.

The Min-Frame Transformation (MFT) is an encoding 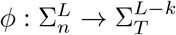 that transforms a DNA sequence *S* of length *L* over the nucleotide alphabet Σ_*n*_ into a representative sequence over an alphabet Σ_*T*_ using a mapping of *k*-mers to characters in Σ_*T*_.

Using a minimizer scheme with ordering 𝒪_*k*_, we first select a single *k*-mer *k*^*\**^ within each window. Without reducing across windows, the selected *k*-mer *k*^*\**^ is then translated via a mapping function *M* : Σ^k^→Σ_*T*_ to produce a representative character for the transformed sequence. This process effectively reduces the window’s information to a character representation of its most minimal *k*-mer.

### ▸Definition 11.

***(MFT Scheme)** An MFT scheme is a tuple (k, w*, 𝒪_*k*_, *and M), which collectively are used to encode a sequence* 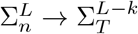.

The combination of (*k, w*, 𝒪_*k*_, and *M*) are collectively referred to for the remainder of this paper as an **MFT scheme**, and modifying each has impact on downstream results (see Appendix for further evaluations of each parameter). Importantly, although we build upon a minimizer sampling scheme in this work, MFT schemes could be defined using any forward sampling scheme that maintains a window guarantee [18, 25]. The primary novelty of the MFT scheme therefore lies in transforming each window into a character over a separate alphabet, rather than simply sampling *k*-mers and their positions.

In **Figure 2**, we show an example MFT scheme, where we utilize *k* = 3, *w* = 5, an alphabetical ordering over 3-mers, and a codon to amino acid mapping table *M*. Despite the presence of a mutation between the two sequences, the resulting transformed sequences are the same.

**Figure 2.**
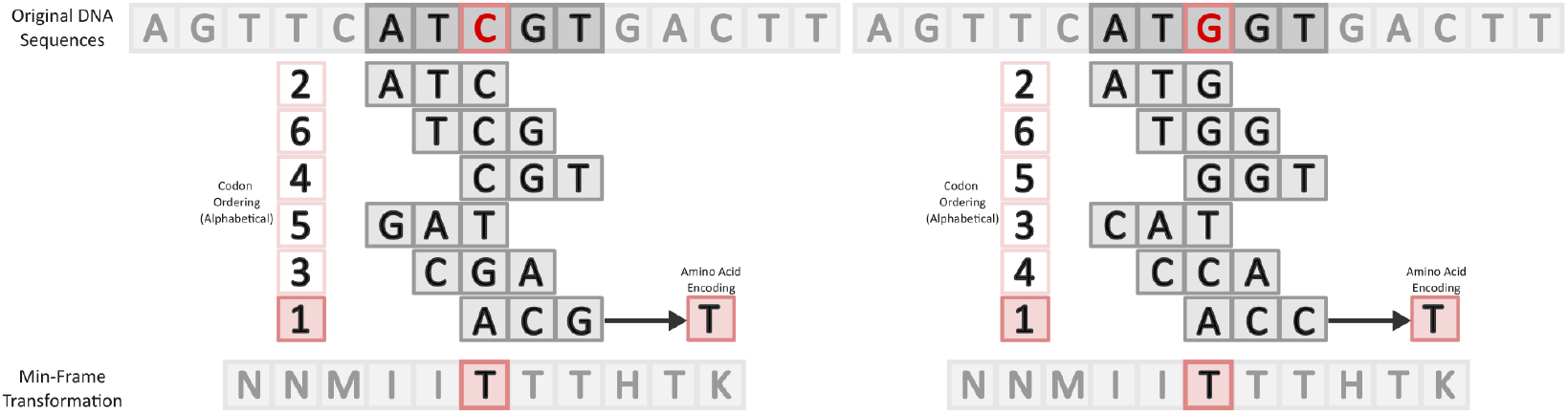
Overview of the Min-Frame Transformation (MFT) on two sequences with a single mutation within them using k=3, w=5, an alphabetical ordering 𝒪_*k*_, and a codon to amino acid mapping function *M*. A window centered on a basepair (in red) is used to enumerate all *k*-mers (both forward and reverse) in that window before using 𝒪_*k*_ to select one and transform it to its final representative character with *M*. The original sequence has a single mutation at the center position, however after the use of the MFT, the resulting transformed sequence no longer has that mutation.

## 4 Results

We focus our primary results on two MFT schemes with *k* = 3, a mapping table *M* derived using a *XX* – matching pattern (alphabet size of 16), a *k*-mer ordering optimized from a random ordering via greedy hill-climbing optimization (see Methods), and window sizes of 5 and 8, respectively. Although numerous other schemes exist, schemes with k=3 are conceptually simple, easy to define, and fast to evaluate exactly given we can quickly enumerate every mutation of every *w*-mer. We include additional evaluations of schemes with k=3 and further discuss this choice in the Appendix.

### 4.1 Real Genomes Experimental Design

The primary goal when identifying maximal unique matches (MUMs) is to recover long, reliable anchors of exact homology across sequences that can guide downstream alignment and identification of SNPs or other genetic variation among the genomes, which enable reconstruction of the evolutionary history of the genome set.

We first evaluated the performance of MFT schemes on real viral genome datasets spanning both low- and high-divergence settings. To assess performance across more divergent genomes, we considered datasets containing multiple genotypes within a species. We then evaluated datasets restricted to a single genotype, representing more homogeneous and less divergent alignments. Specifically, we downloaded 849 complete *Measles morbillivirus* genomes (taxid: 11234) and 686 complete *Human mastadenovirus B* genomes (taxid: 108098), both containing sequences across multiple genotypes, as well as 496 complete *Zaire ebolavirus* genomes (taxid: 186538) and 2,717 complete SARS-CoV-2 genomes (taxid: 2697049) for the single genotype evaluations.

In all cases, we generated five replicates of 200 sequences each, sampling without replacement when possible or otherwise re-shuffling to ensure even representation of sequences across the replicates. For each replicate, we iteratively subsampled genomes to generate nested sets of sizes 100, 75, 25, and 10, such that each smaller set was a subset of the previous larger set, yielding five replicates at each size. The resulting sequence sets from all experiments were transformed using both MFT schemes. Multi-MUMs were then identified in both the transformed and original nucleotide sequences using Mumemto [20], with minimum lengths of 10, 20, and 30. Using the length 20 multi-MUMs, we applied a simplified, modified version of Parsnp2 and custom LCB filtering (see Methods) to generate alignments and extract SNP information.

As primary metrics to evaluate MFT schemes, we evaluated the fraction of the genome aligned and the number of SNP sites recovered. To further characterize the underlying multi-MUM quality, we also assessed multi-MUM coverage and contiguity (see **Appendix Figures 2-3**).

### 4.2 Alignment Coverage and SNP Recovery for Real Viral Genomes

In **Figure 3**, we show alignment coverage and SNP site recovery in sets of *the Measles morbillivirus* and *Human mastadenovirus B* genomes, where both sets of genomes contain multiple divergent genotypes. For both viruses, MFT schemes produced longer alignments with substantially more SNP sites across the board, aside from when only looking at 10 measles genomes. Importantly, the relative alignment length and SNP sites identified increased as the number of sequences grew. For example, with 100 measles genomes, an MFT scheme with *w* = 8 aligned more than 40% of the genome and recovered more than 1,300 SNP sites on average, while the nucleotide-based alignment covered only 10% of the genome and recovered only *∼*300 SNP sites. This corresponds to a 4-fold increase in both the alignment length and recovered SNP sites, which could be critical to reconstruct the evolutionary relationships between the genomes.

**Figure 3.**
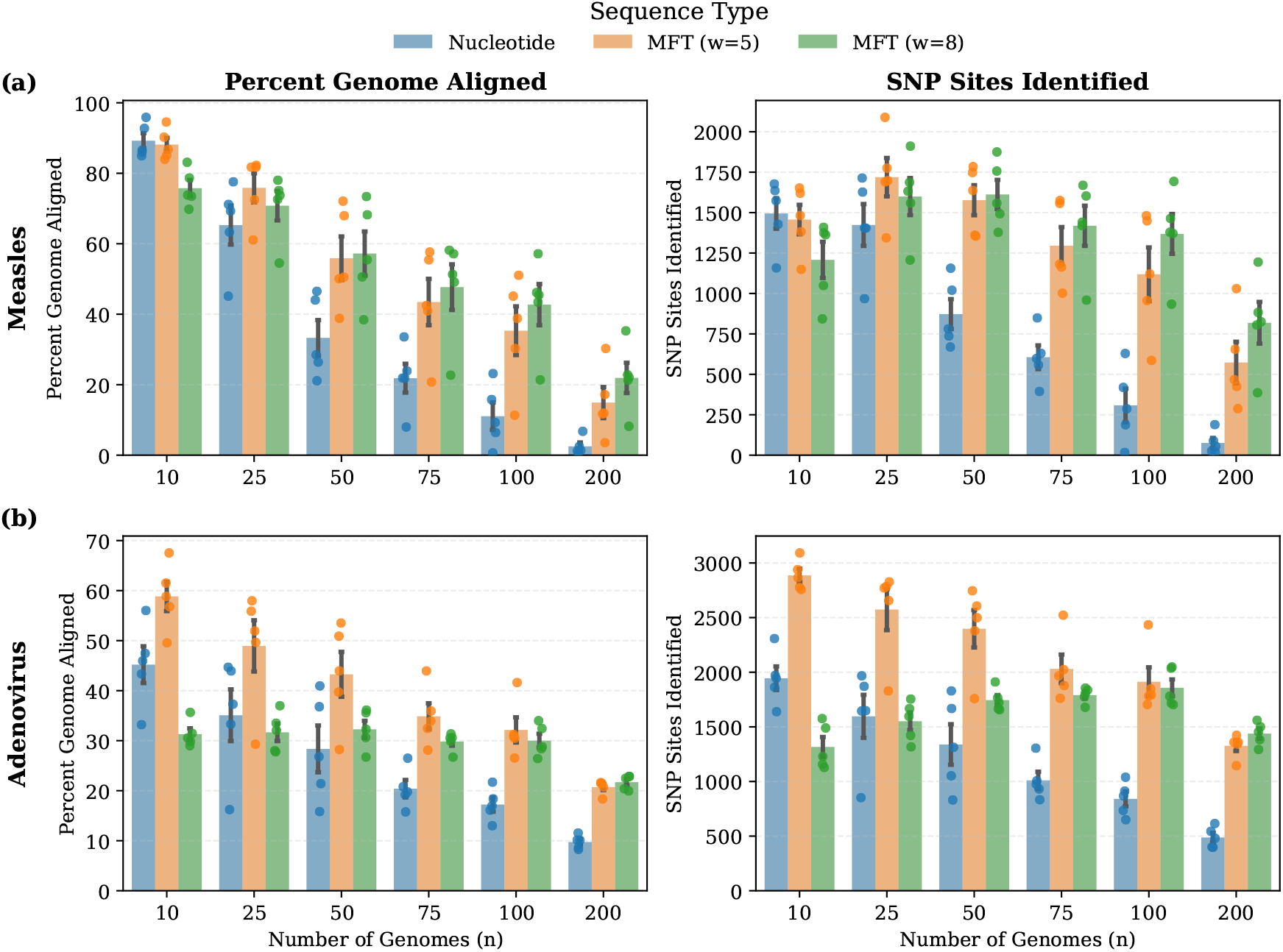
Percent of genome aligned and number of SNP sites recovered for increasing numbers of a) *Measles morbillovirus* and b) *Human mastadenovirus B* genomes. As the number of sequences increases, the use of the MFT leads to consistently longer alignments and more SNP sites than typical nucleotide alignments.

As additional tests on more homogeneous viral populations, we evaluated alignment performance on SARS-CoV-2 genomes sampled across the pandemic and *Zaire ebolavirus* genomes sampled across multiple outbreaks. Unlike the *Measles morbillivirus* and *Human mastadenovirus B* datasets, which span multiple genotypes, these datasets represent more genetically homogeneous populations with lower overall divergence. Accordingly, as shown in **Figure 4**, genome coverage remains stable for both organisms as the number of sequences increases. For example, *Zaire ebolavirus* maintains nearly 70–80% genome coverage even at 200 genomes, while SARS-CoV-2 remains near 100% coverage across all subsets, with both MFT schemes marginally outperforming the nucleotide-based approach at larger set sizes.

**Figure 4.**
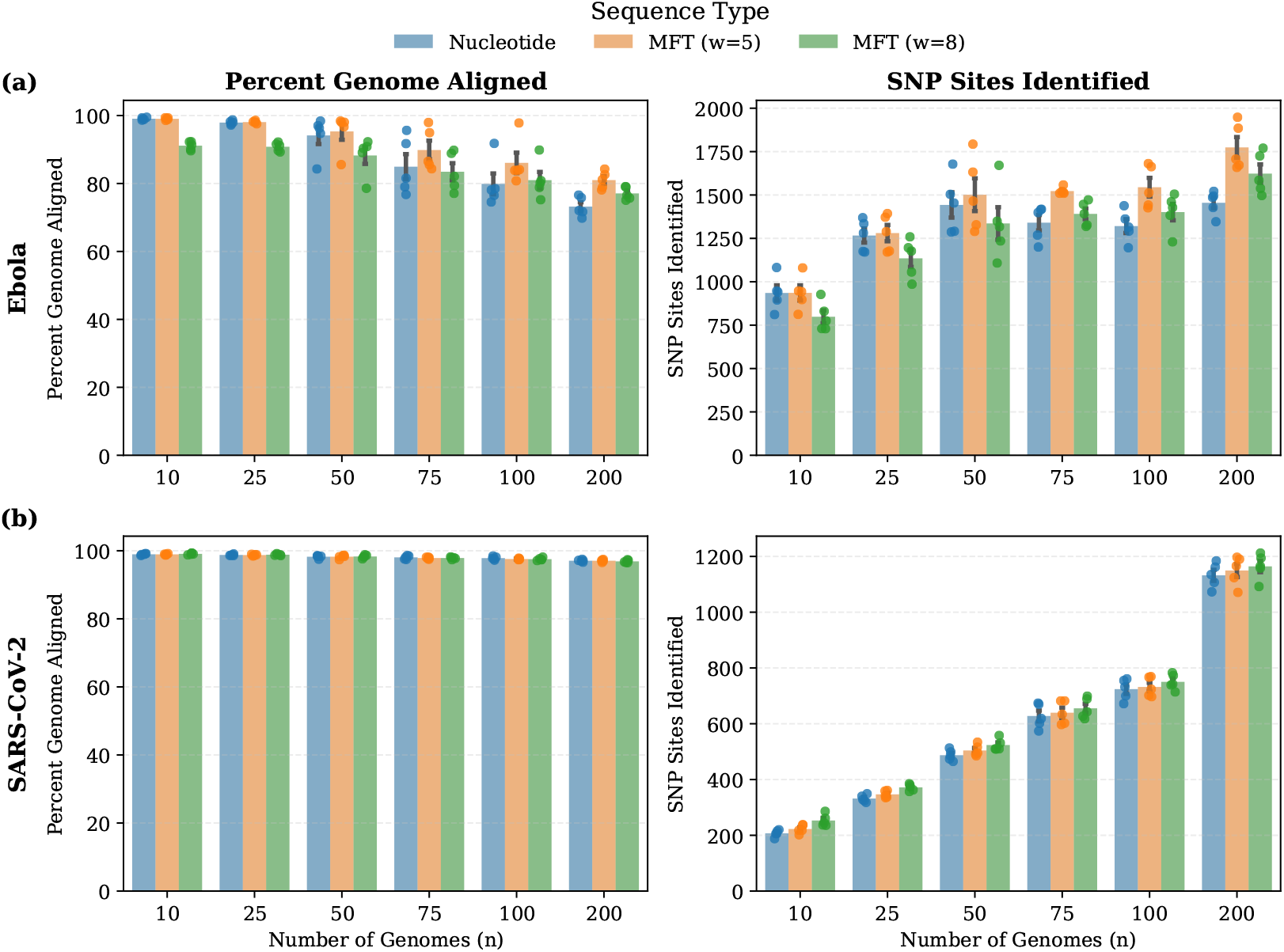
Percent of genome aligned and number of SNP sites recovered for increasing numbers of a) *Zaire ebolavirus* and b) SARS-CoV-2 genomes. MFT schemes (see *w* = 5) achieve modest increases in alignment length and SNP recovery, particularly as the number of genomes increases.

In these settings, improvements from the MFT are more modest. However, despite similar overall alignment coverage, we consistently observe small increases in the number of SNPs recovered across multiple values of *n*. While these gains are smaller than in more divergent datasets, even a limited number of additional SNPs can be critical for applications such as resolving transmission events, where genomes may differ by a handful of nucleotides [26, 27, 28]. Additionally, despite similar final alignment lengths, the underlying multi-MUM structure differs substantially. As seen in **Appendix Figure 2**, MFT schemes produce far less fragmented multi-MUMs, reducing the amount of chaining required to achieve the final alignment. For example, with *w* = 8, MFT-derived multi-MUMs have up to a 50% increase in the multi-MUM N50 compared to those obtained directly from nucleotide sequences.

### 4.3 Simulated Experimental Design

To further evaluate the extent to which MFT improves alignment coverage and SNP recovery, we considered two simulated settings that systematically vary genome divergence and sample size. These simulations provide a highly controlled evaluation of performance by preserving exact positional homology and therefore a known ground-truth relationship between genome sequences. They also represent more challenging conditions than the real viral datasets, since mutations are introduced randomly across the genome without the evolutionary constraints and conserved regions present in real biological sequences [23, 29]. We perform two simulated experiments described here:

1. Random mutations under a star phylogeny. We first generated a random 10,000bp root *−* genome. From this genome, we independently simulated *n* − 1 descendant genomes by randomly introducing single-nucleotide polymorphisms (SNPs) at rate *α*. This setup isolates the theoretical behavior of multi-MUMs under independent divergence. We evaluated a wide range of *α* and *n*, with 10 replicates per parameter setting to account for stochastic variation.
2. Mutations along a viral phylogeny using seq-gen [30]. We constructed a phylogeny from 200 SARS-CoV-2 genomes using Parsnp2 and RAxML [31] and used it to simulate genomes of length 10,000 using a Jukes-Cantor [32] model with a constant rate across all sites (see Appendix for full details). We scaled all branch lengths by a varying multiplicative factor to control the overall evolutionary rate, and sampled leaves to obtain varying values of *n*. Similar to the previous setup, we repeated the simulation over 10 replicates for each set of parameters, but used the same underlying phylogeny.

Similarly to the real viral genome sets, the simulated sequence sets were transformed using both MFT schemes. Multi-MUMs were then identified for both the transformed and original nucleotide sequences with Mumemto [20], using a minimum multi-MUM length of 20 nucleotides. The modified version of Parsnp2 and custom LCB filtering (see Methods) was then used to generate alignments and extract SNP information.

We evaluated performance in the simulated datasets using three metrics. First, we measured alignment coverage, defined as the percent of the genome aligned obtained after running Parsnp2. Second and third, we evaluated SNP precision and recall, which quantify the accuracy of recovered SNPs through the number of true positive and false positive variant calls across methods.

### 4.4 Simulation Results under a Star Phylogeny

In **Figure 5**, we show the results of the first simulated experiment across varied numbers of genomes and SNP mutation rates *α*. First and most importantly, across the two MFT schemes, we observe consistent increases in alignment coverage, with *w* = 8 producing the most pronounced increase (**Figure 5a**). In some configurations of n and *α*, this difference is particularly significant. For instance, with n=10, *α* = 0.02, the alignment coverage jumps from around 40% with the nucleotide-based multi-MUMs to upwards of 60% for a MFT with *w* = 8, realizing a near 50% increase in the alignment length.

**Figure 5.**
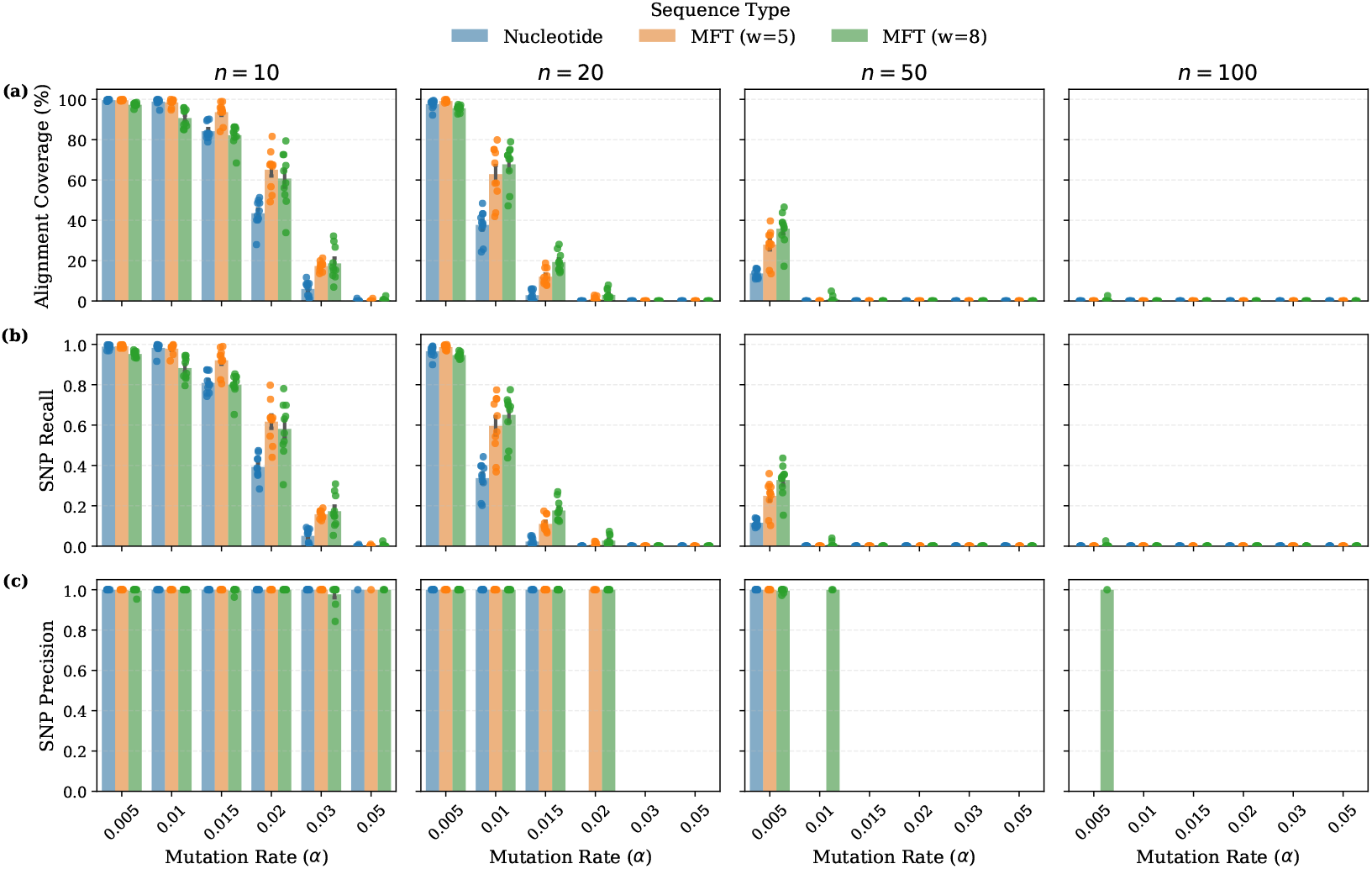
Alignment coverage (a), SNP recall (b), and SNP precision (c) as functions of the mutation rate *α* and number of sequences. The MFT schemes consistently yield longer alignment lengths (a) and higher SNP recall (b), with gains becoming more pronounced as the mutation rate increases. SNP precision (c) remains nearly perfect across all settings when SNPs are found, with only a small number of false positives observed for the MFT scheme with w=8. SNP precision is only shown when at least one SNP is identified.

As a result of the increased alignment coverage, we also observed a corresponding increase in the number of truth SNPs recovered, with SNP recall improving over 0.2 in some cases (**Figure 5b**). Importantly, very few false positive SNPs were identified (**Figure 5c**), and these were limited to a small number of cases for the MFT scheme with *w* = 8, primarily when the number of genomes was small. Consequently, SNP precision remained high whenever SNPs were recovered, typically exceeding 0.95 and remaining at 1 in most settings.

### 4.5 Results on Simulated Genomes Derived from a Real Phylogeny

In the second experiment, we evaluated a simulated setting in which genomes evolved along a known phylogeny. Unlike the random star-phylogeny simulations, this allows mutations to be inherited across related genomes, producing a more biologically realistic correlation structure between sequences. In this setting, we again observed consistent improvement in alignment coverage and SNP recall for MFT schemes, particularly as the branch length multiplier increased (**Figure 6**). In some cases, such as *n* = 25 with a branch length multiplier of 0.75, the MFT schemes resulted in 20% more aligned bases and 0.2 higher SNP recall than the nucleotide-based approach. SNP precision again remained near 1 across nearly all settings when any SNPs were identified, with only a small number of false positive SNPs observed for the MFT scheme with *w* = 8.

**Figure 6.**
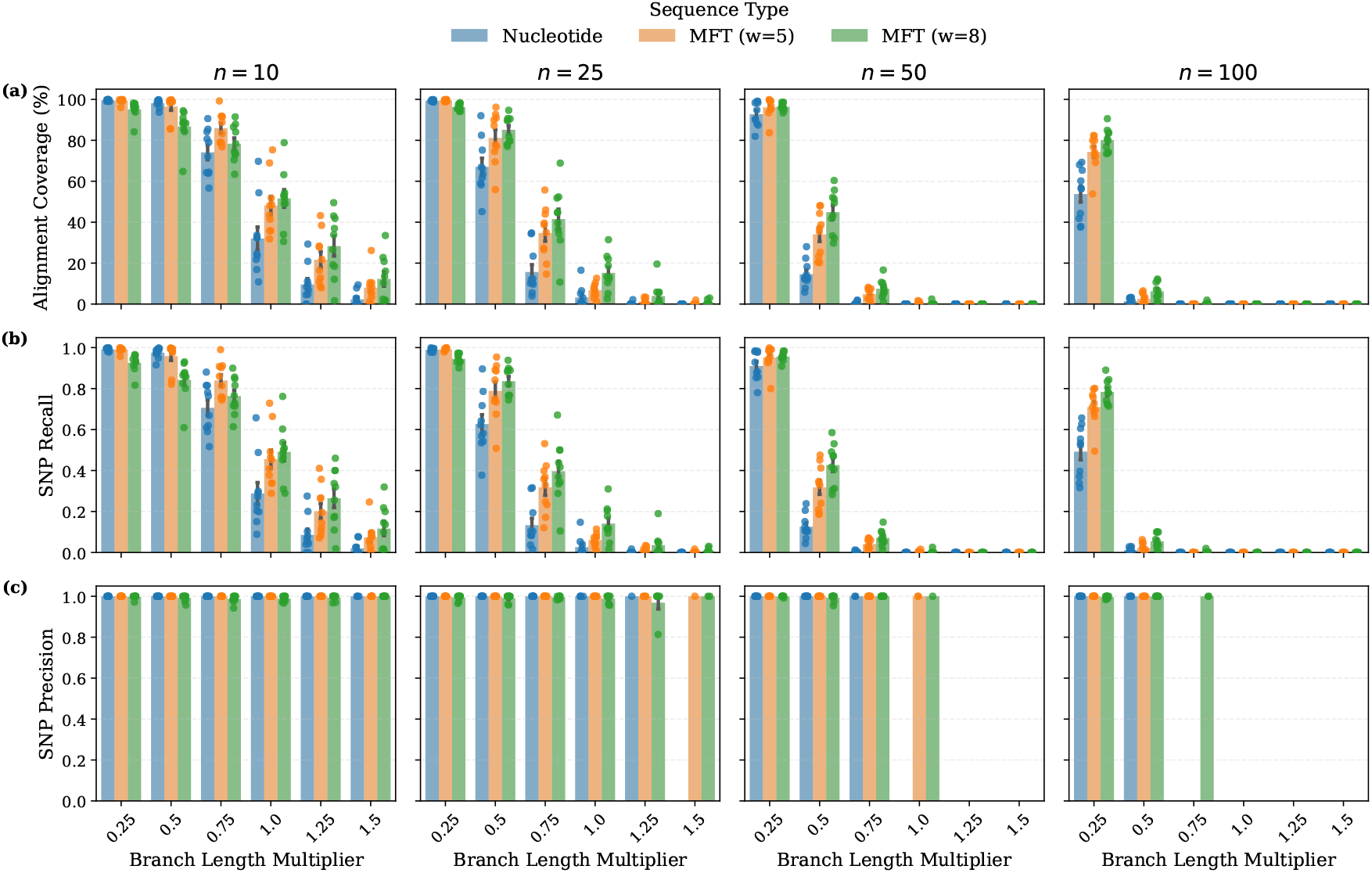
Alignment coverage (a), SNP recall (b), and SNP precision (c) as functions of the branch length multiplier and the number of simulated sequences derived from a SARS-CoV-2 phylogeny under a Jukes-Cantor model. The MFT schemes consistently yield longer alignment lengths (a) and higher SNP recall (b), with gains becoming more pronounced as the mutation rate increases. SNP precision (c) remains nearly perfect across all settings when SNPs are found, with only a small number of false positives observed for the MFT scheme with w=8. SNP precision is only shown when at least one SNP is identified.

Interestingly, the alignment coverage in these simulations contrasts sharply with the real SARS-CoV-2 datasets shown in **Figure 3b**, where nucleotide alignments achieve near-complete genome coverage. This discrepancy suggests that the simulation protocol we followed (Jukes-Cantor site evolution and constant rates across all sites) produce substantially more challenging alignment conditions than real viral evolution, even when mutations occur along a phylogeny. In particular, because all sites evolve under the same rate, there are no highly conserved regions, which is a very different pattern than what is seen in real viral genomes [24, 29, 33]. As such, these simulations should primarily be interpreted as controlled stress tests of alignment robustness under highly randomized mutation processes.

## 5 Discussion

In this work, we introduce the Min-Frame Transformation (MFT) as a fast preprocessing step that transforms DNA sequences into an alternative alphabet to improve alignment via multiple maximal unique matches (multi-MUMs). Across both simulated and real datasets, MFT sequences result in longer alignments with greater SNP site recovery than nucleotide-based approaches, driven by the identification of longer, more contiguous multi-MUMs.

Conceptually, the MFT provides a sensitivity boost analogous to protein-level matching while retaining the simplicity and efficiency of nucleotide multi-MUM–based alignment. Even when improvements in alignment length are modest, the additional recovered SNPs can be critical for downstream tasks such as phylogenetic inference, transmission reconstruction, and marker discovery, reinforcing the practical importance of these gains.

From a theoretical perspective, we show that MFT schemes can achieve up to 90% masking of single SNPs within a window *w*. However, this gain in masking rate is associated with a tradeoff of reduced sequence entropy, which can diminish the availability of short unique sub-strings and negatively impact multi-MUM detection (see **Appendix Tables 1-3**). The primary limitation of the MFT is the trade-off between mutation robustness and sequence uniqueness, where sub-strings that are unique in nucleotide space may no longer remain unique after transformation. Depending on the MFT scheme, this can cause some multi-MUMs to collapse or disappear entirely, particularly in larger collections or longer genomes where there is more chance for similar sub-strings. In practice, several strategies could help mitigate this effect. For example, one could utilize subset multi-MUMs that span most, but not all, sequences while remaining collinear, or incorporate multi-MEMs when anchored by surrounding multi-MUMs. Additionally, recursive multi-MUM discovery, as implemented in Parsnp2, could recover matches within subregions where the uniqueness criterion is more likely to hold. Even without additional algorithmic modifications, the gains from longer and more stable anchors across divergent sequences may outweigh the loss of shorter matches, particularly from the perspective of structural genomics.

Outside of the decrease in sequence uniqueness, there are other properties of the transformation that can make things more challenging for downstream multiple genome alignment. Because we select from both forward and reverse *k*-mers within each window, MFT sequences are effectively canonicalized. For non-collinear genomes with more structural rearrangements such as bacteria, this will necessitate the development of additional algorithms to handle such cases. Alternatively, the MFT could be adjusted to only consider a single strand, but this may necessitate other parameter adjustments given the number of *k*-mers in each window is cut in half. The combination of the decrease in uniqueness with longer sequences and canonicalization of the sequences are the primary motivation for why we focus on viral genomes in this work, as those challenges are not relevant.

Lastly, although we show two MFT schemes perform well in this viral context, performance varies across datasets. For example, the *w* = 5 scheme performed best on the real viral datasets, whereas the *w* = 8 scheme often achieved larger gains in simulated settings. This suggests that MFT schemes may currently depend strongly on the underlying evolutionary model and motivates several directions for future work to improve MFT schemes more generally. First, while we implement and describe MFT schemes using minimizers, alternative sampling schemes such as syncmers [25] may further improve the ability to mask variation, given syncmers offer higher *k*-mer conservation than minimizers [25, 34]. Second, our SNP masking objective and optimization procedure are relatively simple. More sophisticated models of *k*-mer statistics [35] could yield more robust objective functions that could lead to improved MFT outcomes. Similarly, evaluating the theoretical limit of masking is also of interest, analogous to bounds on sampling density [19, 36, 37]. Finally, investigating indel-aware MFT encoding schemes may further enhance performance.

Overall, the Min-Frame Transformation already provides substantial improvements in alignment quality with minimal computational overhead, and more broadly suggests that alphabet transformations are a promising direction for improving exact-match–based comparative genomics.

## 6 Additional Methods

### 6.1 Defining Mapping of *k*-mers to a Transformed Alphabet

Given a *k*-mer size k, we must define a mapping 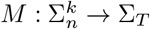that assigns each *k*-mer over the nucleotide alphabet Σ_*n*_ to a character in a transformed alphabet Σ_*T*_. One of the key ideas in the Min-Frame Transformation is to map similar *k*-mers (by edit distance or some other metric) to the same character, similar to numerous other approaches [38, 39, 40, 41]. The primary reasons for the mapping are two fold: 1) Mapping *k*-mers to an ASCII alphabet enables the re-use of MUM finding software. 2) Mapping multiple *k*-mers to the same character enables additional robustness against mutations within a window than simply relying on *k*-mer conservation from a given *k*-mer sampling scheme. There are several ways to define mappings, and we do not comprehensively explore all of them in this work. For mappings that are not pre-defined (e.g., codon-to-amino-acid mappings for *k* = 3), we construct mappings using spaced-seed pattern matching [14, 42, 43]. For example, with *k* = 3, the pattern “XX-” (also defined elsewhere as 110) maps all 3-mers sharing the same first two nucleotides to the same character. The weight *m* of a spaced-seed pattern is defined as the number of match positions and directly determines the size of the transformed alphabet, |Σ_*T*_| = 4^*m*^. Also of note is that these are *balanced* in that they map the exact same number of *k*-mers to each character (4^*k*−*m*^).

Due to this exponential growth, we limit the weight of the spaced seed pattern to 3 or less in mft-tools, as beyond that exceeds the number of machine-readable ASCII characters (95 total). However, there may be more complex ways to encode patterns in the future that enable heavier weighted patterns.

### 6.2 SNP Masking Rate of MFT Schemes

To evaluate the robustness of various MFT schemes, we developed a basic framework to quantify the SNP masking rate, which measures the probability that a single nucleotide polymorphism (SNP) within a sequence window does not alter the resulting MFT character. From here on, we utilize *R*_*m*_ for the SNP masking rate. For a given window size *w*, we iterate over the set of all possible *w*-mers 𝒲, where |𝒲| = 4^*m*^. Then, for each sequence 𝒮 ∈ 𝒲, we first compute the initial transformed character *v* = *ϕ*(*S*) using the given MFT scheme and then systematically test the outcome of the transformation *ϕ* on all 3*w* sequences with a single point mutation. *R*_*m*_ is then mathematically defined as follows:

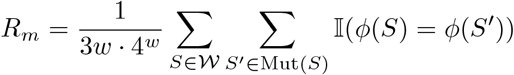

In this expression, Mut(*S*) represents the set of sequences differing from *S* by exactly one base, and 𝕀(·) denotes an indicator function. A higher *R*_*m*_ signifies a MFT scheme is more effective at masking single nucleotide variants. In addition to the global rate, we also calculate transition specific masking rates (ex. *A*→ *T*, *G* →*C*). For each mutated sequence, we track the nucleotide substitution (*b*_1_ →*b*_2_) ∈ ℬ that occurred, where (*b*_1_,→ *b*_2_) andℬ denotes the set of all possible nucleotide substitutions. This allows us to decompose the global metric into a set of 12 transition-specific rates, defined as:

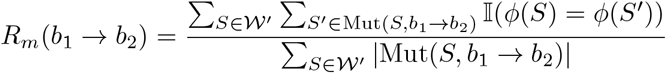

where Mut(*S, b*_1_→*b*_2_) represents the subset of mutations in which a base *b*_1_ at any position in *S* is replaced by a base *b*_2_ in *S*^*′*^. By isolating these rates, we can identify whether ordering schemes exhibit systematic biases, or design schemes tailored to specific organisms by tuning to organism-specific substitution frequencies (see **Appendix Table 3**).

For small *w*, we compute the global and transition specific SNP masking rate value exactly. However, for *w* ≥ 8, due to the number of *k*-mers growing exponentially, we empirically estimate the SNP masking rate by sampling *n* random w-mers *W*. For the remainder of this paper, we subsample *n* = 10000 *w*-mers to generate SNP masking rates for *w* ≥8. In this case, the mapping rate is defined as:

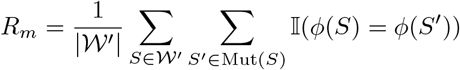

where 𝒲′ ⊆ 𝒲 and |𝒲′| = *n*.

### 6.3 Shannon Entropy of MFT Schemes

While the SNP masking rate measures mutational robustness, it is insufficient on its own for evaluating a mapping scheme. For example, a trivial mapping that assigns every *k*-mer to the same character would achieve a perfect masking rate of *R*_*m*_ = 1.0, but would eliminate all underlying genomic information. Effective MFT schemes must therefore balance mutation robustness with preservation of sequence structure. To quantify this trade-off, we analyze the distribution of transformed values using Shannon entropy. As with the SNP masking rate, we compute the entropy *H*(*X*) by iterating over all *w*-mers (or subset 𝒲^′^for *w* ≥8) and estimating the probability mass function *P* (*v*) for each character *v∈* Σ_*T*_ in the transformed alphabet:

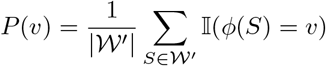

The entropy of the mapping scheme is then defined as:

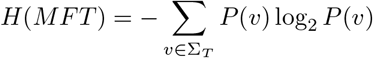

Higher entropy indicates that the ordering and mapping table more effectively utilize the available transformed alphabet, distributing characters more evenly across a broader range of values.

### 6.4 Optimizing orderings

The configuration of the *k*-mer ordering 𝒪_*k*_ is a critical aspect of MFT scheme performance. While an analytically optimal ordering may exist, the combinatorial complexity of the state space makes empirical derivation non-trivial, and something we leave to future work. For now, we define the optimization of 𝒪_*k*_ as a search for an ordering that maximizes the SNP masking rate *R*_*m*_ as the objective function. Given a fixed mapping table and an initial ordering 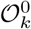, we employ a greedy stochastic hill-climbing algorithm to iteratively refine the ordering 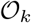, similar to the hill-climbing algorithms used to optimize for minimizer density in [44]. In each subsequent iteration *t*, we generate a candidate ordering 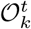 by swapping a random pair of *k*-mers in the current best ordering 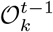. The candidate is accepted if and only if it yields a positive improvement in the objective function:

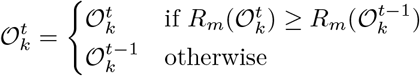

We execute this process for *T* = 2000 iterations, a threshold we observed was near convergence to a local maximum. Additionally, because we can compute masking rates for specific transitions, we also can define a weighted objective function using weights *W* over all 12 possible transitions. This weighted masking rate, *R*_*m,w*_, allows optimization of preferential masking against specific mutations. The weighted objective function is defined as:

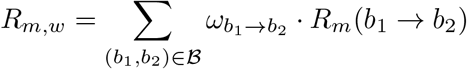

where ℬ is the set of all possible nucleotide substitutions, *R*_*m*_(*b*_1_ → *b*_2_) is the transition-specific masking rate, and 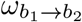 is the weight assigned to that specific mutation. By normalizing these weights such that *ω* = 1, we can tune the ordering 𝒪_*k*_ to be more robust to certain known nucleotide transition frequencies (see **Appendix Table 3**).

### 6.5 MUM inference and genome alignment

Multi-MUM identification was performed with Mumemto v1.4.0 using minimum length thresholds of `-l 10, -l 20`, or `-l 30` as specified. The full `Mumemto` command used for each set of genomes was: `mumemto *.fna -l [min_length] -o [output].` Multiple genome alignment for the real viral genomes was carried out using a modified version of Parsnp2 [7] that allows externally computed MUMs from `Mumemto` to be used directly. Specifically, we added a `--external-mums` flag to use MUMs from `Mumemto`. We also ran Parsnp2 with `--mum-length 10000000` to disable recursive MUM search and ensure only provided MUMs were used, as well as `--no-partition` to align all genomes in a single pass, and `--skip-ani-filter` to retain all genomes from the MUM computation even if divergent. Finally, we set `--min-anchor-length 20` to exclude MUMs shorter than 20bp from serving as alignment anchors. The full command is in the Appendix.

After obtaining the raw alignments from the modified Parsnp2 pipeline, we performed an additional filtering step to retain only locally collinear blocks (LCBs) longer than 50bp that did not break the collinearity of the entire alignment. This reduces spurious alignments that can arise when aligning MFT sequences and provides more accurate estimates of alignment length and SNP recovery. To do this, LCBs were first ordered by length. Starting from the longest LCB, we greedily add each candidate block to the final alignment if all sequences within the block remained collinear with respect to the LCBs already accepted. Final alignment lengths and SNP counts were then calculated from the filtered set of LCBs. Notably, this approach assumes largely conserved genome structure and collinearity, which is generally appropriate for viral genomes outside of known reassortment events [24].

## Acknowledgements

R.D. is supported by a training fellowship from the Gulf Coast Consortia, on the NLM Training Program in Biomedical Informatics & Data Science (T15LM007093)

## 7 Code and Data Availability

All MFT functions are available at https://github.com/treangenlab/mft-tools and downloadable via bioconda (mft-tools). Additional scripts used to generate the results are also available at https://github.com/treangenlab/mft-analyses. The modified version of Parsnp2 that can accept MUMs from Mumemto as input is available at https://github.com/treangenlab/parsnp_external_mums

