## Appendix for "Min-frame transformation enables more sensitive viral genome alignment"

### A Appendix

#### A.1 Comparison of multi-MEMs and multi-MUMs

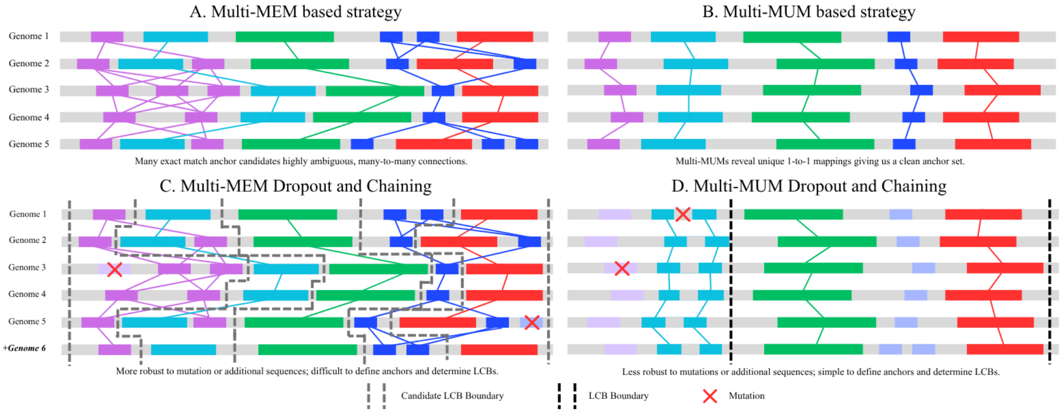


**Appendix Figure 1** Schematic comparison of multi-MEMs (A,C) and multi-MUMs (B,D) for multiple genome alignment seeding. The uniqueness constraint of multi-MUMs provides strict one-to-one anchors for downstream alignment, whereas multi-MEMs produce many-to-many relationships that require substantially more chaining. (C) Multi-MEMs are generally more robust to mutations, but their increased ambiguity makes construction of locally collinear blocks (LCBs) more difficult. (D) In contrast, multi-MUMs can fragment or disappear due to mutations or the introduction of non-unique sequence in additional genomes, but their strict uniqueness makes them substantially easier to chain into LCBs.

#### A.2 Empirical performance of Min-Frame Transformation Schemes

We began by exploring Min-Frame Transformation (MFT) schemes with set parameters $k = 3$ and $w = 5$. We chose these values as a simple baseline that reflects a standard biological six-frame translation when considering a single codon. Additionally, these parameters allow for easily defined mapping tables given the small $k$-mer size, and orderings can be evaluated exactly and quickly due to the small window size, which enables rapid optimization. Finally, this configuration allows us to include known unbalanced mappings in our comparison, such as the standard codon-to-amino-acid translation.

To enable this comparison, we defined three mapping tables to project 3-mers onto a fixed alphabet: two balanced schemes, $X--$ and $XX-$, which map to alphabets of size 4 and 16 respectively (where X denotes a match and $-$ a variable position), and a standard codon translation table as an unbalanced biological reference. For each mapping, we assessed three classes of orderings: a naive alphabetical ordering, a random order, and a final optimized order generated by subjecting the random order to our greedy hill-climbing optimization process. Due to the stochasticity of the random and optimized orderings, we repeated these processes over 100 seeds, and reported the 10-90th percentiles instead of a single point metric.

Looking across the board in **Appendix Table 1**, all MFT schemes achieved masking rates above 35%, suggesting that even for naive random or alphabetical orderings, a significant portion of SNPs can still be masked using MFT schemes. However, the resulting SNP masking rates and entropy values vary widely across schemes. Looking more closely at individual maps, maps that reduce $k$-mers to smaller alphabets ($X--$, e.g.) tend to have larger SNP masking rates, but lower entropy than maps with larger alphabets ($XX-$ and

**Appendix Table 1** SNP masking rates and Shannon entropy for $k = 3$, $w = 5$ across different orderings and mappings

| $k$ | $w$ | $\mathcal{O}_k$ | $M$ | **Mapping Type** | **SNP Masking Rate** | **Shannon Entropy** |
|---|---|---|---|---|---|---|
| 3 | 5 | Alphabetical | $X--$ | Balanced | 77.50% | 1.42 |
| 3 | 5 | Alphabetical | $XX-$ | Balanced | 51.56% | 3.38 |
| 3 | 5 | Alphabetical | Codon | Unbalanced | 45.67% | 3.66 |
| 3 | 5 | Random | $X--$ | Balanced | 54.46—57.85% | 1.93—1.99 |
| 3 | 5 | Random | $XX-$ | Balanced | 38.27—41.40% | 3.74—3.89 |
| 3 | 5 | Random | Codon | Unbalanced | 35.76—38.49% | 3.86—4.11 |
| 3 | 5 | Optimized | $X--$ | Balanced | 75.13—77.50% | 1.42—1.49 |
| 3 | 5 | Optimized | $XX-$ | Balanced | 54.15—56.34% | 3.07—3.19 |
| 3 | 5 | Optimized | Codon | Unbalanced | 51.24—54.42% | 3.01—3.27 |

) *Note:* ($k$, $w$, $\mathcal{O}_k$, $M$) represent the MFT scheme used. Mapping types are balanced when the same number of $k$-mers are mapped to each character in the transformed alphabet. When Masking rate and Shannon entropy are given as a range, the values represent the 10th and 90th percentile over 100 replicates.

codon). This makes sense as with more $k$-mers pointing to the same character, you expect more variation to be masked, while also reducing the character variation of the resulting sequence. On the other hand, the $XX-$ (alphabet size 16) and codon (alphabet size 20) maps yielded slightly lower SNP masking rates, but higher entropies in comparison.

Furthermore, the choice of ordering $\mathcal{O}_k$ plays a critical role in performance. While random orderings still provided reasonable masking rates ranging from $35.76 - 57.85\%$, the masking rates were lower than both alphabetical and optimized versions. The hill-climbing optimization performed directly on the random orderings improved the masking rate across all mapping types. For instance, for the mapping type $XX-$, the SNP masking rates jumped from 38.27–41.40% with the random orderings to 54.15–56.34% with the optimized. However, again, the tradeoff to increase masking rates comes with an associated decrease in Shannon entropy, dropping from 3.74–3.89 to 3.01–3.27. Interestingly, the Shannon entropy of random DNA sequences is 2.0, so the transformed sequences actually can increase in entropy due to the increased alphabet size, despite mapping many $k$-mers to the same character.

We next evaluated the performance of the optimized orderings when the window size was larger. Specifically, we evaluated $w = 8$ and $w = 11$. This allows for 12 and 18 3-mers, respectively, to be evaluated, which gives more opportunities for 3-mers to be chosen from unmutated regions. Given the results shown in Table 1, we focused on the optimized orderings of each mapping type, given that the masking rates were highest.

As shown in **Appendix Table 2**, increasing the window size $w$ enables a clear increase in SNP masking rate across all mapping types. For example, in the $X--$ map, the masking rate climbs from a baseline of 54.15–56.34% at $w = 5$ to 74.71–77.89% at $w = 11$, representing a near 20% increase. However, again accompanying these increased masking rates is a corresponding collapse in sequence entropy. For example, the $XX-$ map sees its entropy drop from roughly 3.07–3.19 for $w = 5$ to 1.91–2.17 for $w = 11$. Intuitively, a broader window context enhances the probability that the ordering $\mathcal{O}$ selects a conserved $k$-mer that avoids local SNPs. However, this also increases the likelihood of the same high-ranked $k$-mers being chosen more often and across adjacent windows, which leads to reduced entropy. The overall result for large window sizes is a highly redundant, lower-entropy representative sequences that are robust to variation but contain less unique information. Given the entropy of optimized schemes with $w = 11$ falls below 2 in many cases, we decided to only include $w = 5$

**Appendix Table 2** SNP masking rates and Shannon entropy for optimized MFT schemes across increasing window sizes

| $k$ | $w$ | $\mathcal{O}$ | $M$ | **Mapping Type** | **SNP Masking Rate** | **Shannon Entropy** |
|---|---|---|---|---|---|---|
| 3 | 5 | Optimized | $X--$ | Balanced | 75.13—77.50% | 1.42—1.49 |
| 3 | 8 | Optimized | $X--$ | Balanced | 83.85—89.21% | 0.79—1.06 |
| 3 | 11 | Optimized | $X--$ | Balanced | 89.45—94.84% | 0.44—0.79 |
| 3 | 5 | Optimized | $XX-$ | Balanced | 54.15—56.34% | 3.07—3.19 |
| 3 | 8 | Optimized | $XX-$ | Balanced | 67.27—70.22% | 2.30—2.52 |
| 3 | 11 | Optimized | $XX-$ | Balanced | 74.71—77.89% | 1.91—2.17 |
| 3 | 5 | Optimized | Codon | Unbalanced | 51.24—54.42% | 3.01—3.27 |
| 3 | 8 | Optimized | Codon | Unbalanced | 66.42—70.23% | 2.12—2.57 |
| 3 | 11 | Optimized | Codon | Unbalanced | 74.15—78.97% | 1.66—2.22 |

) *Note:* ($k$, $w$, $\mathcal{O}_k$, $M$) represent the MFT scheme used. Mapping types are balanced when the same number of $k$-mers are mapped to each character in the transformed alphabet. Values for the SNP masking rate and Shannon entropy are given as the 10th and 90th percentile over 100 replicates.

and $w = 8$ in the main text.

Additionally of note, the codon and $XX-$ mappings maintain comparable performance throughout the scaling process, despite different mapping tables and alphabet sizes.

#### A.3 Transition-specific optimizations

After optimizing to the global SNP masking rate, we wanted to investigate the possibility of identifying orderings that mask certain mutation types better than others and match known variation patterns in organisms of interest. As a proof of concept, we tuned the optimization such that certain transition pairs of the same two nucleotides (i.e, $A \to C$, $C \to A$) were weighted 4x as much in the optimization score as all other mutation types. We repeated this for a handful of nucleotide pairs (C/T, C/G, and A/G) and then report the global SNP masking rates, the global Shannon entropy, and then target specific masking rates, as well as non-target masking rates (where targets are defined as the transition pairs heavily weighted in the optimization).

As seen in **Appendix Table 3**, the target transitions masking rates are significantly higher than the global masking rates, while non-target masking rates range more widely from below the global rate to slightly above. This suggests that there exist orderings that can be tuned towards certain mutational patterns. Importantly, the upper ranges of the global masking rates are similar to those when the weights are even, suggesting that you can achieve global masking rates that are competitive, even if certain mutation types are preferentially masked. However, the lower end of the global masking rates are lower, suggesting that there may be fewer orderings within the space of all possible orderings that exist for these weights.

#### A.4 MUM Statistics for Viral Organism Experiments

In **Appendix Figures 2-3**, we present raw multi-MUM statistics for the same measles and SARS-CoV-2 virus subsets presented in the main text, filtering across three minimum length thresholds. Specifically, we analyze the multi-MUM coverage, which represents the length of multi-MUMs identified, as well as the multi-MUM N50. Analogous to its use genome assembly, the multi-MUM N50 is defined as the largest value $\ell$ such that all multi-MUMs of length $\ell$ or greater collectively account for at least 50% of the total multi-MUM

**Appendix Table 3** SNP masking performance across weighted optimizations ($k$=3, $w$=5, ordering='XX-'). Target columns (T1, T2) correspond to mutation types emphasized by each optimization scheme; non-target rates summarize all remaining mutation types.

| Weight | Global SNP Masking Rate | Global Entropy | Targets Masking Rates T1 | Targets Masking Rates T2 | Non-Target Masking Rates |
|---|---|---|---|---|---|
| | | | - | - | |
| Even | 54.15—56.34% | 3.07—3.19 | - | - | - |
| | | | $C \to T$ | $T \to C$ | |
| C/T | 51.52—55.56% | 3.10—3.32 | 63.94—73.36% | 63.94—73.36% | 44.20—57.03% |
| | | | $C \to G$ | $G \to C$ | |
| C/G | 51.40—54.14% | 3.11-3.32 | 63.44—73.14% | 63.44—73.14% | 44.91—57.12% |
| | | | $A \to G$ | $G \to A$ | |
| A/G | 50.88—55.58% | 3.11-3.36 | 64.57—73.29% | 64.57—73.29% | 43.89—56.57% |

sequence length. Higher N50 values indicate longer, less fragmented multi-MUMs, which are preferable for downstream alignment. In most cases, MFT-transformed sequences outperform nucleotide-based approaches in both total multi-MUM length and N50, indicating that the gains in alignment coverage and SNP recovery are primarily driven by an increase in longer multi-MUMs spanning larger portions of the genome.

An exception arises at larger window sizes ($w = 11$ for instance) and higher sequence counts ($n > 50$) when allowing multi-MUMs of length 10, where we observe a slight reduction in alignment length. This is primarily due to increased collisions among short multi-MUMs, where sequences that are unique in nucleotide space can become non-unique after transformation, thus reducing the number of valid short matches. This effect becomes more pronounced as the number of sequences (and thus the probability of collision in at least one sequence) increases. However, when restricting to multi-MUMs longer than 30, MFT multi-MUMs yield substantially greater coverage via longer matches, which generally provide higher confidence of homology than shorter multi-MUMs.

#### A.5 Simulated Phylogeny Parameters

We first generated a core genome over 200 SARS-CoV-2 genomes using Parsnp2 v2.1.5 using the following command: `parsnp -g *.fna -r GCA_964157915.1.fna.ref -p 32 -o cov_tree`. The full list of NCBI accessions are available in the mft-analyses Github repository. Within Parsnp2, RAxML v8.2.13 [31] is used to generate the best maximum likelihood tree using the following command: `raxmlHPC-PTHREADS -m GTRCAT -p 12345 -T 16 -s /path/to/output/parsnp.snps.mblocks -w /path/to/raxml_output -n OUTPUT`

Seq-gen v1.3.5 was then used to generate simulated sequences from the best maximum likelihood tree using the following command: `seq-gen -mGTR -l10000 -ofs -s[scale] -z[seed] -y test` across multiple branch length scales and 10 replicates defined by unique seeds.

#### A.6 Modified Parsnp2 Command

We ran the modified version of Parsnp2 with the following command: `./parsnp -d *.fna -o /path/to/output -p 32 -external-mums /path/mumemto.mums -fo -no-partition -skip-ani-filter -min-anchor-length 20 -mum-length 1000000 -skip-phylogeny`

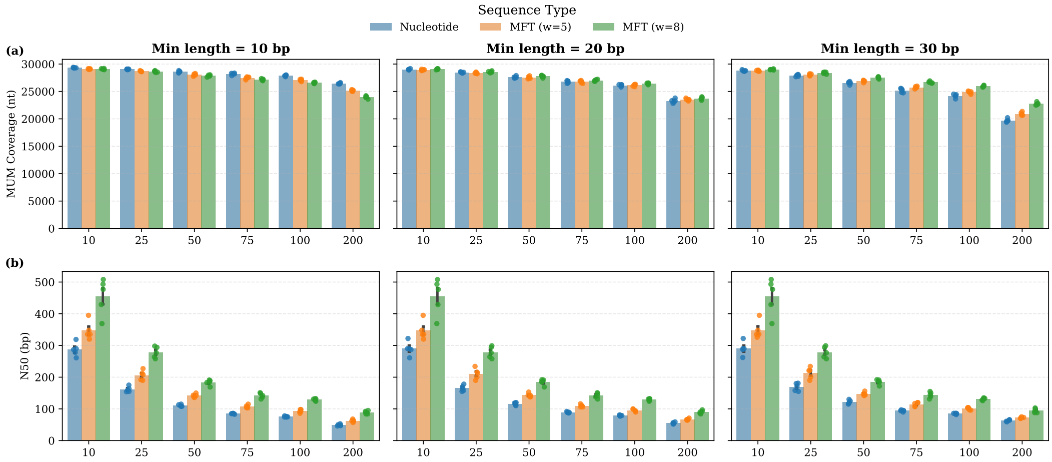


**Appendix Figure 2** Comparison of multi-MUM genome coverage (a) and multi-MUM N50 (b) across increasing sequence counts for SARS-CoV-2 genomes. Multi-MUMs were evaluated under three minimum length thresholds. Nucleotide-based multi-MUMs tend to produce more short, fragmented matches, whereas MFT-based approaches yield longer, more contiguous multi-MUMs.

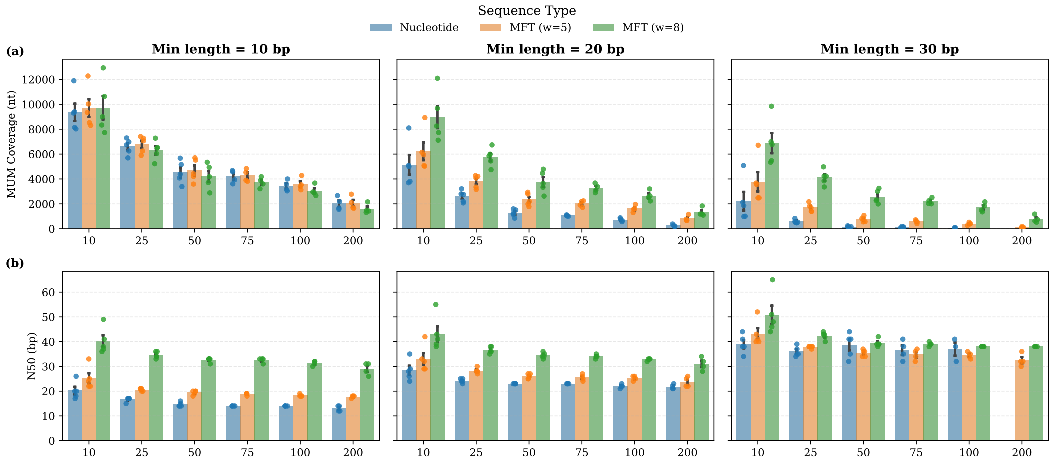


**Appendix Figure 3** Comparison of multi-MUM genome coverage (a) and multi-MUM N50 (b) across increasing sequence counts for Measles genomes. Multi-MUMs were evaluated under three minimum length thresholds. Nucleotide-based multi-MUMs tend to produce more short, fragmented matches, whereas MFT-based approaches yield longer, more contiguous multi-MUMs.
